# Coronavirus M proteins disperse the trans-Golgi network and inhibit anterograde protein trafficking in the secretory pathway

**DOI:** 10.1101/2025.09.05.674545

**Authors:** Taylor Caddell, Rory P. Mulloy, Jennifer A. Corcoran, Eric S. Pringle, Craig McCormick

**Affiliations:** Department of Microbiology & Immunology, Dalhousie University, 5850 College Street, Halifax, NS B3H 4R2, Canada; Department of Microbiology, Immunology and Infectious Diseases, University of Calgary, 2500 University Drive NW, Calgary AB, Canada T2N 1N4; Snyder Institute for Chronic Diseases and Charbonneau Institute of Cancer Research, University of Calgary, 2500 University Drive NW, Calgary AB, Canada T2N 1N4

**Keywords:** Virus, coronavirus, SARS-CoV-2, HCoV-OC43, Membrane, Spike, unfolded protein response, ATF6, STING, Rab26

## Abstract

Coronaviruses (CoVs) encode Spike, Membrane (M), and Envelope (E) transmembrane proteins that are translated and processed at the endoplasmic reticulum (ER) and traverse the secretory pathway to converge at sites of virus assembly. Three transmembrane ER resident proteins, activating transcription factor 6 (ATF6), inositol-requiring enzyme 1 (IRE1), and PKR-like endoplasmic reticulum kinase (PERK), sense the accumulation of unfolded proteins in the ER and initiate the unfolded protein response (UPR) to increase ER protein folding capacity. We observed UPR modulation by numerous CoV proteins, including Spike, which broadly activated all three arms of the UPR. By contrast, M selectively inhibited the ATF6 arm of the UPR, either when it was activated by CoV proteins like Spike, or when it was activated by chemical stimuli of ER stress; however, M was unable to inhibit Spike-mediated activation of IRE1 or PERK. ATF6 inhibition was conserved amongst all human CoV M proteins. Amongst the UPR sensors, ATF6 has a unique activation mechanism whereby ER stress triggers translocation to the Golgi where ATF6 is processed by resident proteases to release the ATF6-N bZIP transcription factor. Because M had no effect on the function of the ATF6-N transcription factor, we reasoned that it may act upstream by inhibiting ATF6 trafficking in the secretory pathway. Indeed, we observed that ectopically expressed M inhibited several processes that require ER-to-Golgi transport, including sterol regulatory element binding protein-2 (SREBP2)-mediated activation of sterol responses and stimulator of interferon response cGAMP interactor 1 (STING1)-mediated activation of interferon responses. M also inhibited the secretion of a soluble Gaussia luciferase reporter protein. Using a Retention Using Selective Hooks (RUSH) cargo sorting assay, we observed that M accumulated in the cis-Golgi and inhibited further anterograde transport of a transmembrane reporter protein beyond this compartment, while dispersing the *trans*-Golgi network (TGN). We also observed a conserved TGN dispersal phenotype in cells infected with SARS-CoV-2, hCoV-OC43, or hCoV-229E. We determined that M is present in both detergent resistant and detergent soluble membranes and that M increased cholesterol abundance at the *cis*-Golgi. Together, these observations suggest that CoV M proteins disrupt the TGN and impede normal anterograde traffic in the canonical secretory pathway, potentially by increasing cholesterol levels at the *cis*-Golgi . Because CoV egress does not require the TGN, this mechanism could allow the virus to selectively interfere with host responses to infection without impeding egress of nascent virions.

**IMPORTANCE:** Many viruses encode proteins that limit host antiviral responses. Coronaviruses encode a remarkably diverse array of proteins that antagonize host responses at discrete steps including the detection of viral products, signal transduction, host mRNA processing and nuclear export, and protein synthesis. Here, we describe a new form of viral antagonism of host antiviral responses by remodelling the secretory pathway, dispersing a distal portion of the Golgi network and causing accumulation of proteins early in the Golgi. This is achieved by an abundant viral structural transmembrane glycoprotein M, which is best known for its role as the central player in the assembly of new viruses. This newly discovered function of M allows it to limit host antiviral responses that depend on trafficking in the late secretory pathway, while maintaining its role in virus assembly in the early secretory pathway.

## INTRODUCTION

Coronaviruses (CoVs) are enveloped viruses with large positive sense single stranded RNA ((+)ssRNA) genomes. Upon entry, the viral genome is released into the host cell cytoplasm and is immediately translated at the endoplasmic reticulum (ER) into long polyproteins that are processed by two viral cysteine proteases, generating a collection of non-structural proteins (NSPs)^1^. Among these, NSP3, NSP4, and NSP6 are ER-localized transmembrane proteins required for formation of ER-derived double-membrane replication organelles (ROs)^2–4^, the site of viral RNA (vRNA) synthesis^5^; this vRNA synthesis is mediated by the RNA-dependent RNA polymerase (RdRp) holoenzyme complex, comprised of the NSP12 polymerase and NSP7/NSP8 co-factors that operate in the RO interior^6^. This intense ER-associated viral protein synthesis and sculpting of ER membranes into ROs are events that could be reasonably expected to perturb ER proteostasis.

Three ER-localized transmembrane proteins are activated in response to the accumulation of unfolded proteins in the ER lumen: activating transcription factor 6 (ATF6), inositol-requiring enzyme 1 (IRE1), and PKR-like endoplasmic reticulum kinase (PERK). Activation of these proteins leads to the generation of basic leucine zipper (bZIP) transcription factors (TFs) that orchestrate the unfolded protein response (UPR) by transactivating genes involved in increasing ER protein folding and processing capacity^7,8^. ER stress triggers the formation of disulfide bonded ATF6 homodimers that migrate from the ER to the Golgi where they are cleaved by resident proteases, liberating the cytosolic amino-terminal ATF6 fragment comprising the ATF6-N bZIP TF^9–11^. By contrast, in response to ER stress, IRE1 generates a bZIP TF via a cytosolic endonuclease domain that cleaves *X-box binding protein 1 (XBP1)* mRNA at 2 sites, followed by ligation of products by the cytosolic RtcB RNA ligase, removing a small intron and shifting an open reading frame to generate the XBP1s bZIP TF^12–15^. Finally, ER stress triggers PERK-mediated eIF2*α* phosphorylation^16^, limiting guanine nucleotide exchange on eIF2^17,18^, and favouring uORF skipping on the *ATF4* mRNA required for synthesis of the ATF4 bZIP TF^19^. Accumulating evidence indicates complex interactions between CoVs and the UPR, whereby infection activates UPR sensors^20–25^, but downstream UPR transcriptional responses are quite limited, as in the example of mouse hepatitis virus (MHV) infection, which triggered *XBP1* mRNA splicing and ATF6 cleavage, but transcription of UPR-responsive genes was attenuated^26^. While our understanding of viral factors that regulate the UPR remains incomplete, several viral UPR agonists have been identified in the literature, including Spike proteins from diverse CoVs^22,27,28^, as well as SARS-CoV transmembrane proteins NSP6^29^, ORF3a^30^, and ORF8ab^31^ proteins.

CoV transmembrane structural proteins, Spike, Membrane (M), and Envelope (E), are translated at the ER and traverse the secretory pathway for further modifications. Spike is a large type I transmembrane protein that receives co-translational N-glycan modifications at the ER that support proper folding, trimerization, and ER export. Spike must be activated by site-specific proteolysis to generate the S1 attachment subunit and an S2 fusion subunit to create infectious CoV virions^32,33^; this processing can be mediated by Golgi-resident proteases, or uncleaved Spike can be incorporated into viral particles and post-egress cleavage can be mediated by cell surface or lysosomal proteases^32,33^. E is a small type I transmembrane protein that forms pentamers and functions as a viroporin, increasing the lumenal pH in the Golgi and protecting Spike from excessive proteolysis in this compartment^34^. M, the most abundant protein in the viral envelope, is a multi-spanning membrane protein, with a small lumenal amino-terminus, three transmembrane domains, and a large cytoplasmic carboxy-terminal domain known as the endodomain^35^. Ectopically expressed M displays steady-state accumulation in the Golgi^36,37^, but it can also be retrieved to earlier compartments, including the ER. This bidirectional traffic is key to its role in virus assembly and budding at the ERGIC, allowing M to capture other structural proteins and redirect them to assembly sites, even after they have trafficked to the *trans* Golgi. These interactions are largely governed by the endodomain, which interacts with the other structural proteins (Spike, E and N)^38–40^. Furthermore, M-M interactions are thought to drive membrane curvature and exclusion of host proteins during envelope formation^41^. Despite being represented in sub-stoichiometric amounts, E assists M in controlling the trafficking and processing of Spike, and as assembly takes place, E acts as an enhancer of budding. Together, these structural proteins, along with nucleoprotein (N), coordinate assembly of CoV virions at the ER-Golgi Intermediate Compartment (ERGIC), followed by lysosomal egress^42^ and release.

In this study we aimed to identify SARS-CoV-2 proteins that modulate the UPR. We identified the M protein as an inhibitor of the ATF6 branch of the UPR, as well as additional signalling proteins like STING that require movement through the secretory pathway to elicit downstream transcriptional responses. We discovered that M accumulates at the *cis*-Golgi, impeding protein trafficking in the canonical secretory pathway, which correlates with *trans-*Golgi network (TGN) dispersal. Because CoV egress does not require the TGN, this mechanism could allow the virus to selectively interfere with host responses to infection without impeding egress of nascent virions. Furthermore, we discovered M causes accumulation of cholesterol at the *cis*-Golgi, suggesting M leads to altered membrane composition at this site. Changes in sterol and lipid abundances in the Golgi can result in perturbations in protein trafficking and cellular secretion, providing a potential mechanism through which M broadly inhibits host intracellular trafficking of proteins requiring ER-Golgi translocation.

## RESULTS

### SARS-CoV-2 proteins modulate ATF6-dependent transcription

ATF6 is a monomeric type II transmembrane ER-resident protein that responds to proteotoxic stress by forming disulfide-linked homodimers that traffic to the Golgi upon activation. At the Golgi ATF6 is cleaved by resident proteases, liberating the cytosolic amino-terminal bZIP transcription factor called ATF6-N that migrates to the nucleus and binds ER stress response elements (ERSEs) to transactivate genes encoding chaperones, foldases, and lipogenesis factors. Thus, we reasoned that ATF6 is an excellent candidate protein to probe the effects of viruses on ER proteostasis and trafficking in the early secretory pathway. We screened the Krogan library of plasmids encoding SARS-CoV-2 open reading frames (ORFs) fused to 2X-Strep tags for the ability to affect ATF6-N-dependent transcriptional responses^43^. HEK 293T cells were transfected with this collection of plasmids, along with a plasmid encoding a firefly luciferase gene driven by two ERSE elements, and a second plasmid bearing a constitutively expressed Renilla luciferase gene that served as a normalization control. We found that four SARS-CoV-2 proteins significantly increased ATF6-N-dependent firefly luciferase activity, including Spike, NSP4, ORF8 and ORF10 (**Figure 1A**); amongst these, we noted that Spike and NSP4 are well documented ER-localized transmembrane glycoproteins, with NSP4 being involved in sculpting ER membrane into viral ROs, and Spike, which had been shown to activate the UPR, but not ATF6 specifically, in previous studies^22,27,28^. We confirmed Spike-mediated ATF6 activation using an untagged Spike protein, which displayed even more robust induction of ERSE-luciferase activity, whereas other SARS-CoV-2 structural proteins E, M, and N, had no significant effect (**Figure 1B**). We used this untagged Spike construct as an agonist in our counter-screen for SARS-CoV-2 proteins that inhibit ATF6-N-dependent ERSE-luciferase activity; we identified seven such proteins, including NSP1, NSP5, NSP14, ORF3a, ORF6, ORF7b, and M (**Figure 1C**). Among these, NSP1 and NSP14 yielded expected results as they are host shutoff proteins and should therefore inhibit expression of luciferase from the reporter constructs, with NSP14 stimulating production of N7-methyl-GTP that inhibits the nuclear cap-binding complex, hindering subsequent pre-mRNA splicing, 3’end processing and export^44^, whereas NSP1 inhibits cap-dependent translation^45–49^. We selected M as our inhibitor of interest because of its well-documented control of Spike trafficking and virus assembly at the ERGIC^40^, and as it has been shown other CoVs activate the UPR combined with the fact M is a conserved viral protein across all CoVs, made it an intriguing target to explore. We demonstrated that the untagged M construct also significantly decreases Spike-mediated ERSE-luciferase activity (**Figure 1D**). The untagged E protein moderately inhibited Spike-mediated ERSE-luciferase activity in this assay, even though the 2XStrep-tagged E protein did not (**Figures 1C, 1D**). This discrepancy in results between the 2xStrep-tagged E and untagged E protein prompted us to carry forward our experiments with the untagged constructs when possible.

**Figure 1.**
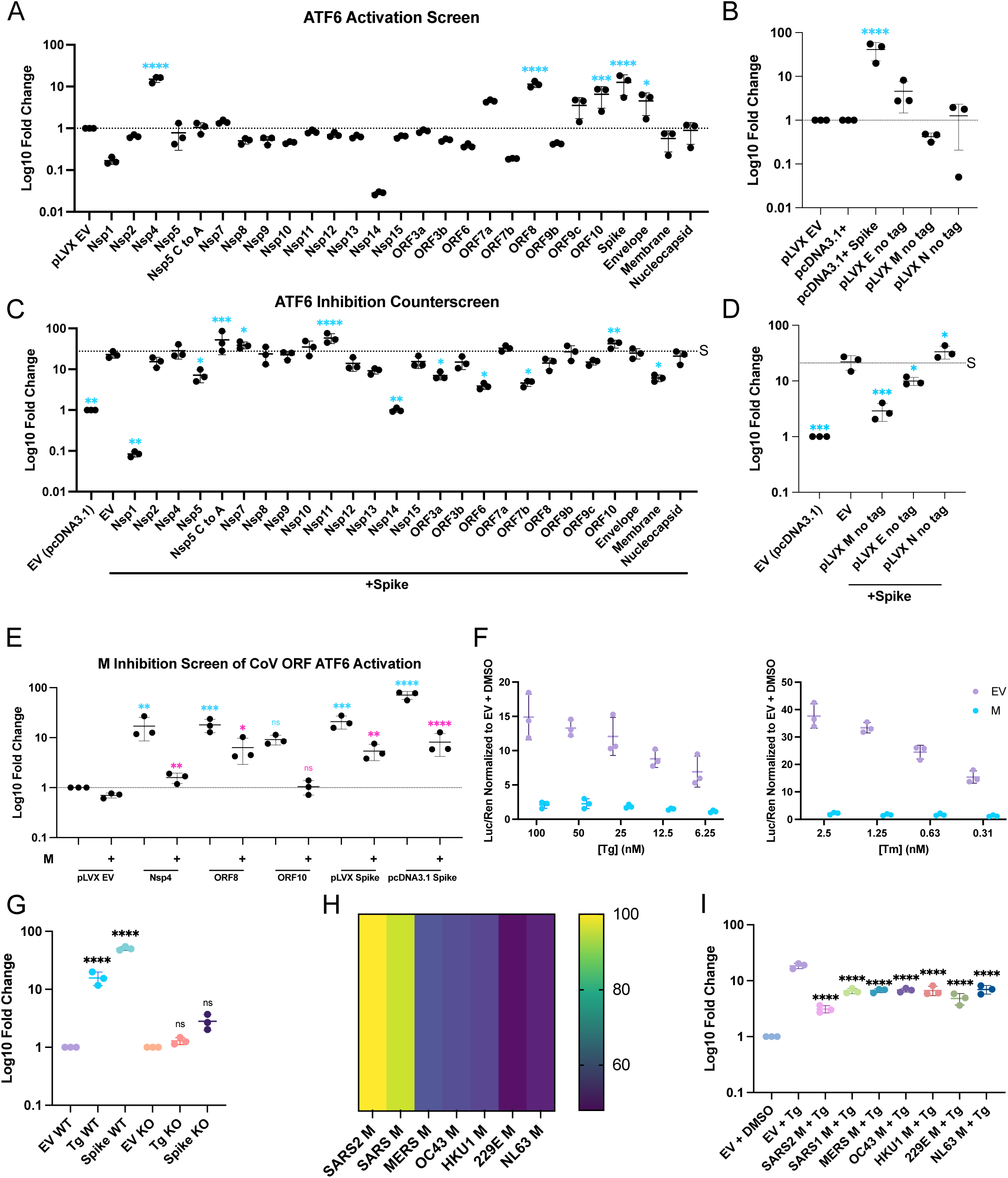
Ectopic expression of SARS-CoV-2 proteins modulate ATF6-dependent transcription. (A-E) SARS-CoV-2 ORF screen for ATF6 activators and inhibitors. HEK293T cells were co-transfected with pLVX vectors encoding viral ORFs plasmids with ESRE-*luc* and CMV-Ren reporters for 24h prior to harvest for luciferase assay. **(A)** Krogran SARS-CoV-2 ORF library where all ORFs encode a C-terminal 2x-Strep tag. **(B)** As in A with untagged SARS-CoV-2 structural proteins. **(C)** A counter-screen for inhibitors was conducted as in **(A)** with the addition of Spike as an ATF6 agonist. **(D)** As in **(B)** with the addition of Spike as an ATF6 agonist. **(E)** as in **(D)** with the addition of M as an ATF6 antagonist as indicated. Blue asterisk indicate significance relative to EV, pink stars indicate significance of M co-transfection relative to paired viral ORF alone **(F)** HEK293T cells were transfected with M then treated as indicated with thapsigargin (Tg) or tunicamycin (Tm) the following day for 24h prior to harvest at 48h. (G) Wild-type HEK293T or ATF6-KO cells were transfected with Spike or treated with Tg. Heat map depicting degree of amino acid similarity between hCoV M proteins. **(I)** as in **(F)** where cells were transfected with M proteins from seven coronavirus known to infect humans or EV. (n = 3 ±SEM, statistical significance was determined by one-way ANOVA with Fisher’s LSD test. *, adjusted P value < 0.05; ***, adj. P < 0.0002; and ****, adj. P < 0.00001 relative to EV.)

We considered the possibility that M’s inhibition of Spike-mediated ATF6 activation was due to well documented physical interactions between Spike and M cytoplasmic tails^40^. To determine whether M possesses broader ATF6 inhibiting activity, we tested it in combination with other ATF6-activating SARS-CoV-2 proteins. We observed once again that NSP4 and ORF8 significantly increased ERSE-luciferase activity, and that M suppressed this phenotype (**Figure 1E**). We also tested whether M could inhibit ATF6 activation by chemical agonists. As expected, treatment of cells with Thapsigargin (Tg), which triggers ER stress by inhibiting the sarcoplasmic/endoplasmic reticulum calcium ATPase 2 (SERCA2)^50^, and Tunicamycin (Tm), which causes ER stress by inhibiting N-linked glycosylation leading to improper protein folding^51^, both triggered ATF6 activation in a dose-dependent manner (**Figure 1F**). M significantly inhibited ATF6 activation at all doses tested for both agonists. We confirmed the specificity of our ATF6 reporter assay by demonstrating that neither Spike nor Tg could stimulate ERSE-luciferase activity in ATF6 knock-out (KO) cells (**Figure 1G, Supplementary Figure 1**). Together, these findings demonstrate that M broadly inhibits ATF6 activation.

As befits its role as a key structural protein that drives CoV assembly, M is quite conserved amongst the hCoVs, with SARS-CoV M most closely related to SARS-CoV-2 M (**Figure 1H**). To determine whether M inhibition of ATF6 activation was a conserved activity of hCoV M proteins, we tested M proteins from all hCoVs for the ability to inhibit Tg-induced ATF6 activation. Remarkably, M proteins from all seven hCoVs potently inhibited ATF6 activation, suggesting that conserved features of these proteins must govern the inhibitory phenotype (**Figure 1I**).

### M inhibits the ATF6 branch but not the PERK or IRE1 branches of the UPR

The UPR consists of three different branches with distinct signaling mechanisms. ATF6 activation triggers anterograde transport and cleavage by Golgi-resident proteases to generate the ATF6-N bZIP transcription factor, whereas IRE1 and PERK generate bZIP transcription factors from other transcripts via cytoplasmic mRNA splicing (XBP1s) and uORF skipping (ATF4) mechanisms, respectively (**Figure 2A**). As our ATF6 reporter assays revealed differential control of ATF6 transcriptional activity by the SARS-CoV-2 structural proteins Spike, E, and M, we embarked on a broader investigation of their effects on all three arms of the UPR. We ectopically expressed Spike, M, and E in HEK293T cells and harvested lysates for immunoblotting. As expected, treatment of cells with Tg activated all three UPR branches, with (i) PERK activation indicated by the presence of a slow-migrating phosphorylated PERK (indicated with a *) and increased production of the ATF4 target CHOP, (ii) IRE1 activation indicated by accumulation of XBP1s, and (iii) ATF6 activation indicated by increased levels of BiP (**Figure 2B**). Ectopic expression of Spike caused activation of all three branches of the UPR as well, whereas E and M showed a minor increase in CHOP expression levels, with E also leading to accumulation of phosphorylated PERK. Neither E nor M had any appreciable impact on the IRE1 and ATF6 branches in these cells.

**Figure 2.**
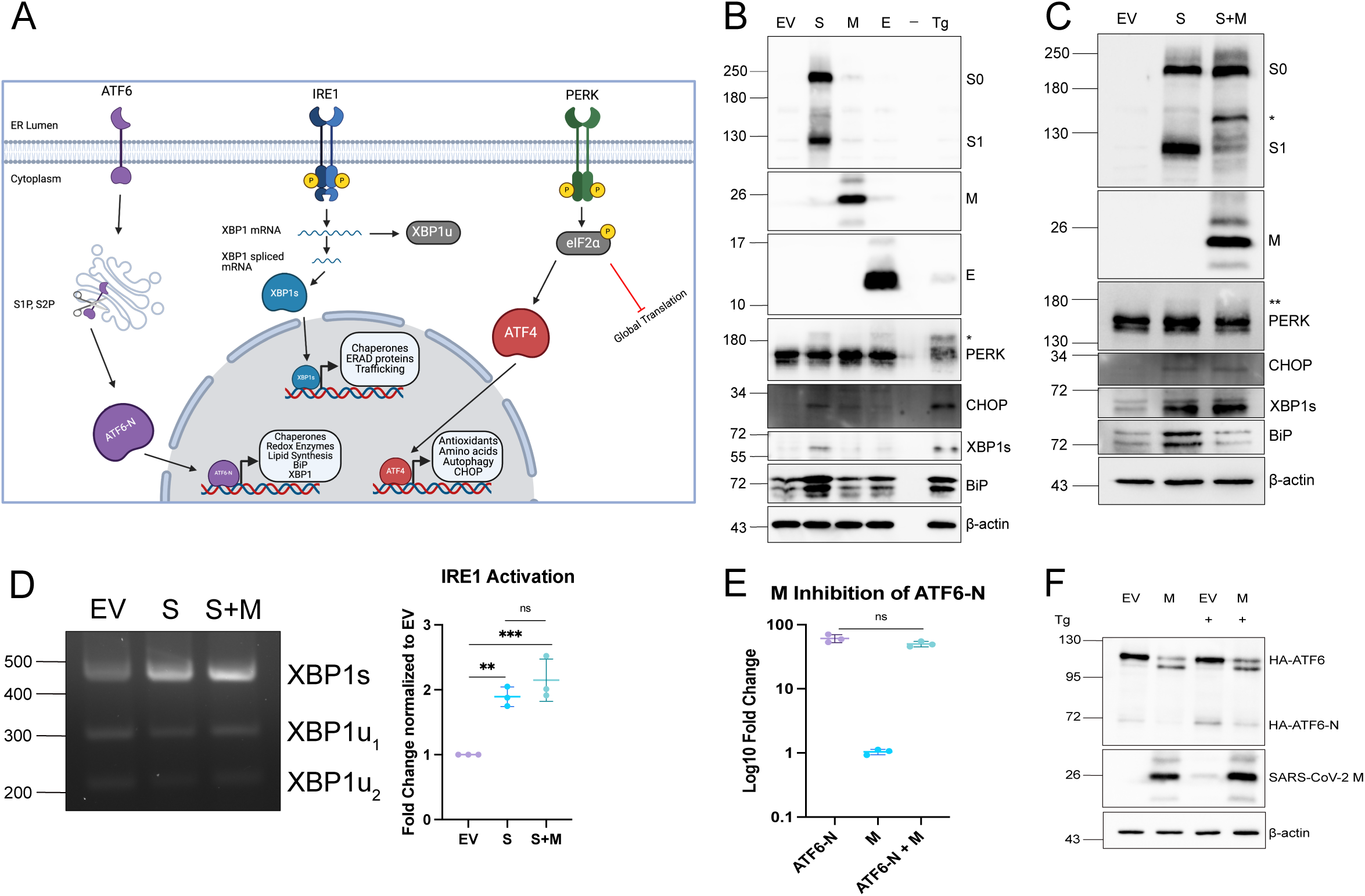
SARS-CoV-2 M protein inhibits ATF6 activation but not PERK or IRE1 activation. **(A)** Diagram of the unfolded protein response (UPR) in brief displaying the intermediate steps that occur for activation and generation of transcription factors that lead to individual transcriptional responses. **(B-C)** HEK293T cells were transfected with SARS-CoV-2 structural proteins as indicated for 24h before lysate was harvested for western blotting. As a positive control untransfected cells were treated with 200 nM thapsigargin (Tg) for 4h prior to harvest. **(D)** as in **(B)** except RNA was harvested for RT-PCR for *Xbp1* mRNA splicing assay. **(Left)** agarose gel; **(Right)** quantification of image by densitometry. **(E)** HEK293T cells were co-transfected with ESRE-*luc*, CMV-Renilla, and HA-ATF6-N with or without M as indicated for 24h prior to harvest for luciferase assay. **(F)** as in **(B)** where cells were transfected with HA-ATF6 with either M or EV control. Cells were treated with 200 nM Tg treatment for 4h prior to harvest as indicated. (For blots, representative image of n = 3 is shown. Otherwise n = 3-4 ±SEM, statistical significance was determined by one-way ANOVA with Fisher’s LSD test. ns – not significant; *, adjusted P < 0.05).

As M inhibits Spike-mediated ATF6 activation, we investigated the effect of M on Spike-mediated activation of IRE1 and PERK. We confirmed that M inhibited Spike-mediated ATF6 activation via diminished accumulation of BiP protein compared to control (**Figure 2C**). Conversely, M had no effect on Spike-mediated activation of PERK (PERK phosphorylation and CHOP accumulation), or IRE1 (XBP1s accumulation). Furthermore, M did not inhibit accumulation of spliced XBP1 mRNA, (**Figure 2D**) Thus, while Spike activates all three branches of the UPR, M appears to only limit activation of ATF6.

ATF6 activation differs from PERK and IRE1 activation because it requires ATF6 translocation to the Golgi for cleavage by resident proteases to generate the ATF6-N transcription factor that migrates to the nucleus to transactivate target genes (depicted in **Figure 2A).** We first considered the possibility that M could inhibit the activity of the ATF6-N transcription factor rather than the precursor ATF6 protein. To test this, we transfected ATF6-deficient HEK293T cells with an ATF6-N construct, either alone or in combination with M, and measured ERSE-luciferase activity. Ectopic ATF6-N expression strongly activated the ERSE reporter, and M had no significant effect on this activity (**Figure 2E**). This suggested that M must inhibit ATF6-N at an earlier stage in the ATF6 activation pathway, prior to cleavage at the Golgi. This was confirmed by our observation that M inhibited the cleavage of an HA epitope-tagged ATF6 construct into an HA-ATF6-N product, either in the presence or absence of Tg (**Figure 2F**).

### M inhibits activation of host pathways that require anterograde transport in the secretory pathway

Selective inhibition of the ATF6 arm of the UPR by M prompted us to investigate other host transmembrane proteins that are similarly activated at the ER and require anterograde transport to the Golgi to elicit downstream transcriptional responses. First, we examined sterol regulatory element binding protein-2 (SREBP2) activation because, like ATF6, it is activated in the ER and processed by Golgi resident proteases to release an amino terminal cytosolic portion of the protein comprising the bZIP transcription factor SREBP2-N^52^. Using a reporter construct bearing three sterol response elements (SREs) that direct transcription of a firefly luciferase gene^53^, we tested the effects of M on sterol responses. Sterol depletion of HEK293T cells by (i) serum starvation or (ii) treatment with statins that inhibit *de novo* sterol synthesis via HMG-CoA reductase inhibition, both increased SRE-luciferase activity (**Figure 3A**). By contrast, cells that expressed SARS-CoV-2 M protein were limited to baseline SRE-luciferase activity in response to sterol depletion. We also investigated the function of the cyclic GMP-AMP synthase (cGAS)/stimulator of interferon genes (STING) pathway. At rest, STING is an ER-resident homodimeric transmembrane protein^54^. Binding to cyclic GMP-AMP (cGAMP) triggers a conformational switch in STING homodimers that directs anterograde transport to the Golgi^54,55^, where it is palmitoylated, and then translocated to the TGN where it is phosphorylated by TANK-binding kinase 1 (TBK1)^56^. Phospho-STING forms a signalling platform at the TGN, directing TBK1-mediated phosphorylation of IRF3, triggering IRF3 dimerization and translocation to the nucleus, resulting in a type I interferon response, and TBK1/IkB kinase epsilon (IKK-ε)-mediated phosphorylation of downstream targets, resulting in activation of NF-κB transcriptional responses^57,58^. To test STING pathway function, we ectopically co-transfected HEK293T cells with cGAS and STING constructs, as well as firefly luciferase reporter constructs bearing interferon-stimulated response elements (ISREs) or NF-κB response elements. cGAS/STING co-transfection activated ISRE-luciferase (**Figure 3B**) and NF-κB-luciferase (**Figure 3C**) reporters as expected. By contrast, co-transfection of M with these complexes caused a marked decrease in output from both reporters. Thus, in addition to ATF6, M also inhibits the SREBP2 and cGAS/STING pathways, all of which traverse the Golgi to execute their respective transcription programs. Finally, to determine whether M could affect the trafficking of a soluble lumenal protein in the secretory pathway, we tested a secreted Gaussia luciferase construct. Cells were transfected with the secreted Gaussia luciferase construct followed by harvest of cell lysates and cell supernatants for luciferase assays. Brefeldin A, a drug that inhibits the GBF1 guanine nucleotide exchange factor for Arf1-GTPase, thereby inhibiting vesicular transport between the ER to the Golgi and causing fusion of ER and Golgi membranes^59–61^, dramatically reduced secretion of the Gaussia enzyme (**Figure 3D**). M also greatly diminished Gaussia secretion. Thus, M causes broad defects in anterograde transport in the secretory pathway for transmembrane and soluble lumenal protein cargo.

**Figure 3.**
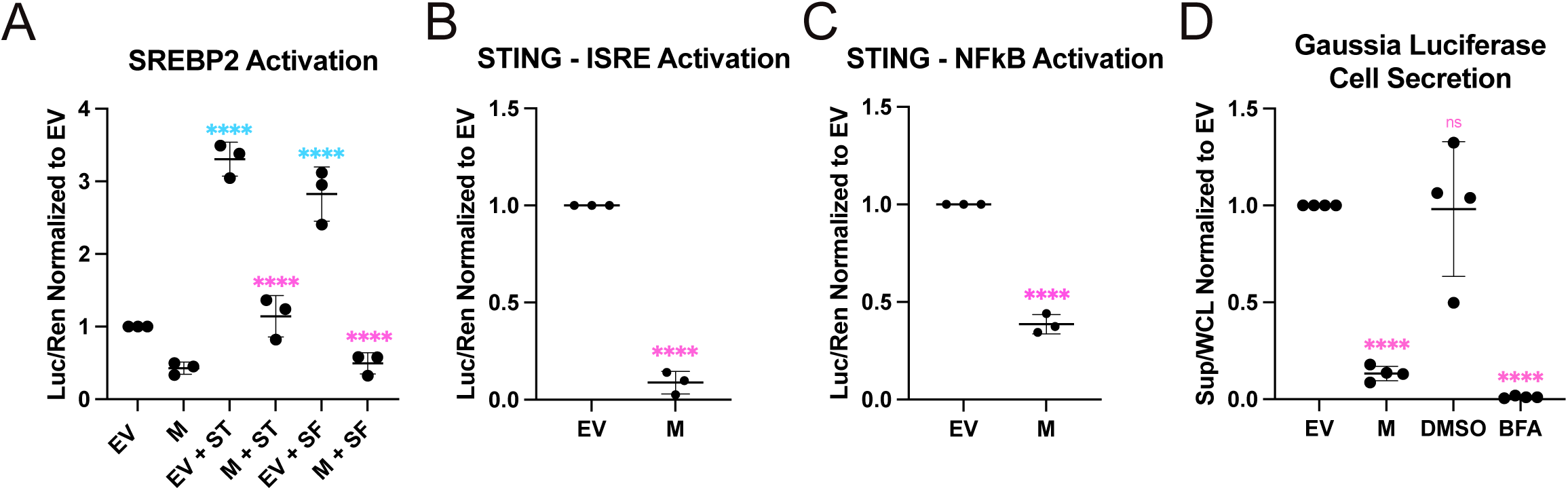
SARS-CoV-2 M broadly inhibits activation of proteins requiring ER-Golgi translocation and cellular secretion. **(A)** HEK293T cells were co-transfected a LDLR-luc, CMV-Renilla, and with M or EV as indicated. The following day cells were treated with 10 µM cerivastatin (ST) or pleased in serum-free media (SF) for 24h prior to harvest for luciferase assay. **(B-C)** HEK293 cells were co-transfected with an **(B)** ISRE-*luc* or **(C)** NFkB-*luc* with cGAS and STING encoding plasmids (to stimulate ISRE and NFkB activation) and with either M or EV as indicated. Cells were harvested 24h for luciferase assay. (D) HEK293T cells were transfected with secreted *Gaussia* luciferase and EV or M as indicated. 24h after transfection, medium was removed and replaced with fresh media. As a positive control cells were treated with 10 µM of Brefeldin A (BFA) or DMSO vehicle control for 6h as indicated. Both supernatant and cells were then harvested at 30h post-transfection for luciferase assay. (n = 3 ±SEM, statistical significance was determined by one-way ANOVA with Fisher’s LSD test. ns – not significant; *, adjusted P < 0.05; ****, adj. P < 0.00001 relative to EV).

### M localizes to the *cis*-Golgi and disperses the structure of the *trans*-Golgi network

With evidence for M-mediated inhibition of anterograde protein trafficking, we investigated the position of M in the secretory pathway. Consistent with previous observations, we observed that ectopically expressed SARS-CoV-2 M accumulated in the *cis*-Golgi^36,37^, as indicated by co-localization with GM130, a peripheral membrane protein of the *cis* Golgi involved maintaining organelle structure^62^ (**Figure 4A**). By contrast, we did not observe steady-state co-localization between M and ERGIC-localized lectin ERGIC53 in HEK293T cells stably expressing GFP-ERGIC53 (**Figures 4A, 4B**), or with the *trans*-Golgi Network (TGN) marker TGOLN2/TGN46 (**Figure 4B**), supporting predominant accumulation of M in the *cis*-Golgi in these cells. We also observed TGN dispersal and reduced staining for TGOLN2 in cells expressing M, suggesting that this compartment may be disrupted.

**Figure 4.**
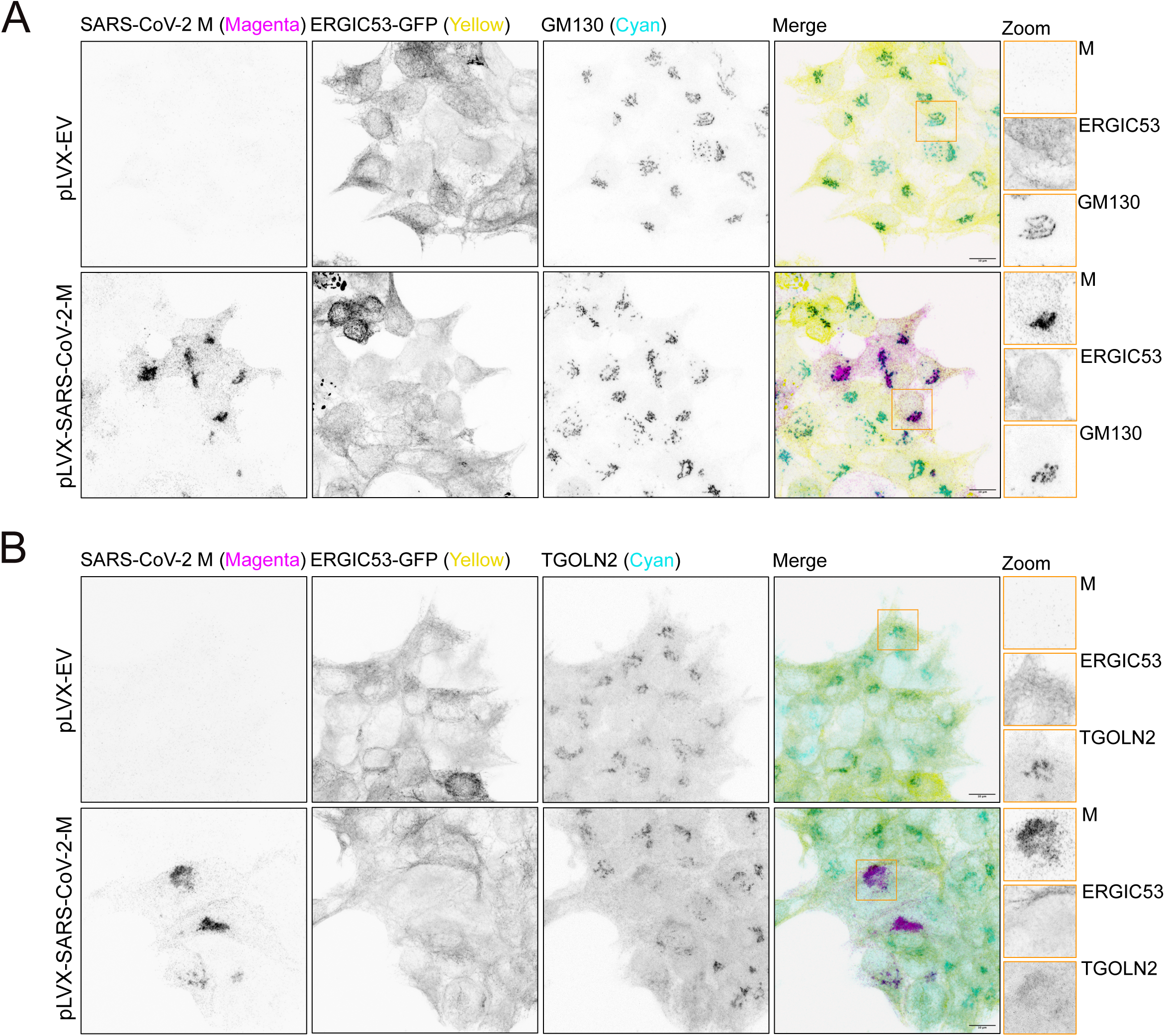
SARS-CoV-2 M localizes to the cis-Golgi and disperses the structure of the *trans*-Golgi network. ERGIC53-GFP-HEK293T cells were transfected with M or EV control and fixed 24h later. Cells were immunostained as indicated with antibodies targeting SARS-CoV-2 M, GM130 (*cis*-Golgi), or TGOLN2 (*trans*-Golgi) and imaged by confocal microscopy. Maximum intensity projections are presented. 100X magnification, scale bar = 10 µm, Orange boxes indicate zoomed field of view. Representative images of three independent experiments.

SARS-CoV-2 and other hCoV infections have been shown to cause Golgi fragmentation and dispersal^63–66^. These studies used TGOLN2/TGN46 as a Golgi marker for dispersal, which we have also found to disperse in the presence of SARS-CoV-2 M; however, as we identified an accumulation of SARS-CoV-2 M at the *cis*-Golgi using the marking GM130, we wanted to further investigate how the *cis*-Golgi and TGN appear during infection. To determine whether CoVs cause *cis*-Golgi dispersal as well as the previously reported TGN dispersal, we infected A549-ACE2 cells with SARS-CoV-2 and immunostained with an anti-M antibody along with markers for *cis*-Golgi (GM130) and TGN (TGOLN2), which revealed that SARS-CoV-2 M accumulates in the *cis*-Golgi during infection, with a fragmented staining pattern for both the *cis*-Golgi and the TGN compared to uninfected cells (**Figure 5A)**. We also observed fragmentation and dispersal for both the *cis*-Golgi and the TGN in hCoV-OC43 (**Figure 5B)** and hCoV-229E (**Figure 5C**) infections, indicating that this phenotype is conserved amongst divergent hCoVs.

**Figure 5.**
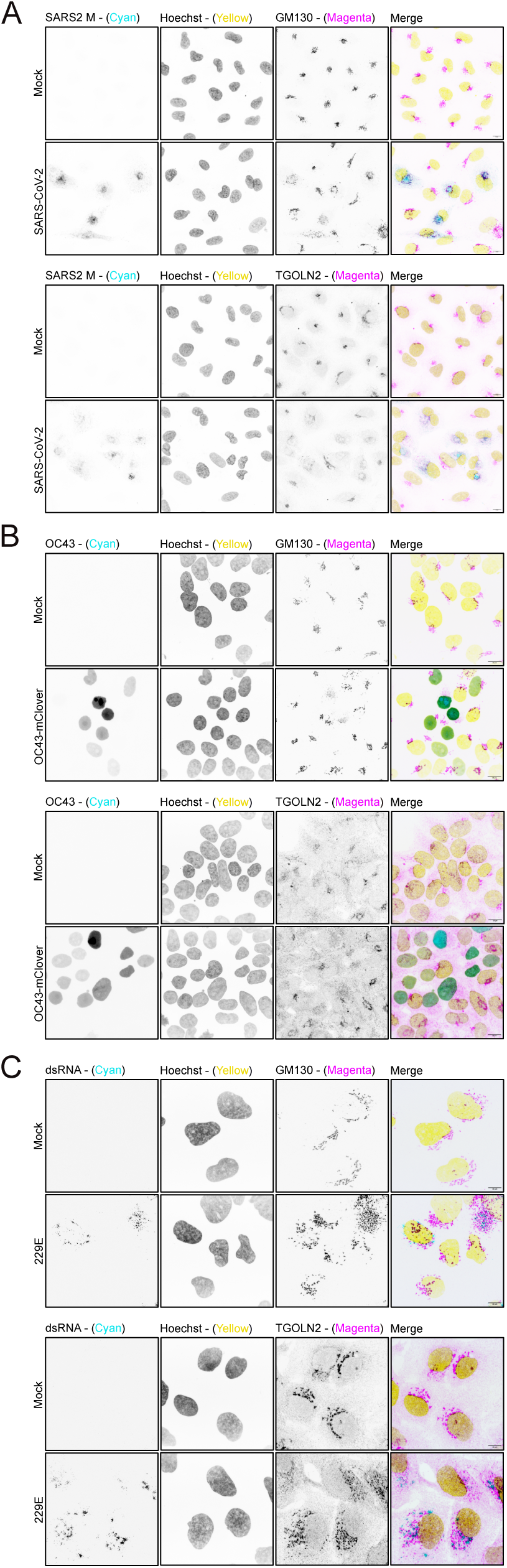
Human coronavirus infections lead to reduced structure and dispersal of both the *cis*-Golgi and *trans*-Golgi network. Confocal immunofluorescence images of GM130 (*cis*-Golgi), or TGOLN2 (*trans*-Golgi) in coronavirus infected cells. **(A)** ACE2-A549 cells were infected with SARS-CoV-2 at an MOI of 1 then fixed at 24hpi. Infected cells were identified with antibodies targeting SARS-CoV-2 M. **(B)** HEK293T cells were infected with hCoV-OC43-mClover at an MOI of 0.05 then fixed at 24hpi. Infected cells express nuclear mClover from a transgene in the recombinant virus. **(C)** Huh7.5 cells were infected with hCoV-229E at an MOI of 0.05 then fixed at 24hpi. Infected cells were identified with the J2 monoclonal antibody that binds dsRNA. Maximum intensity projections are presented. **(A)** 63X, **(B-C)** 100X magnification, scale bar = 10 µm. Representative images of three independent experiments.

### M restricts protein trafficking through the secretory pathway resulting in host protein accumulation at the *cis*-Golgi

We used a Retention Using Selective Hooks (RUSH) cargo sorting assay^67^ to measure the effects of SARS-CoV-2 M protein on dynamic protein trafficking in the secretory pathway in living cells (**Figure 6A**). This assay employs a two-component system comprised of an ER-restricted ‘hook’ construct that includes a streptavidin domain, and a ‘reporter’ construct that includes a streptavidin binding peptide (SBP) fused to a fluorescent protein. As these proteins accumulate in the cell they co-localize in the ER via streptavidin-SBP interactions. The addition of biotin competes with this interaction and releases the SBP-containing reporter construct from the ‘hook’, allowing synchronous trafficking of reporter proteins through the secretory pathway. We used a type II ER-retained Streptavidin-Ii hook protein (this encodes a version of the human invariant chain of MHC that contains an amino-terminal motif for ER retention) and a type II transmembrane tumor necrosis factor (TNF) protein domain fused to SBP and the mApple fluorescent protein. We observed that the reporter construct accumulated in the ER as expected, with punctate structures representing previously described ER exit sites (ERESs) and no signal co-localization with the distal *cis*-Golgi compartment^68^ (**Figure 6B**). After 24 h and 4 h of biotin treatment, the reporter protein formed large puncta that partially co-localized with the *cis*-Golgi marker, displaying proper translocation and trafficking through the secretory pathway, as well as more distal puncta closer to the cell periphery (indicated by black arrows), indicative of vesicular trafficking toward the cell surface^67^. In cells expressing SARS-CoV-2 M, the reporter protein had a largely similar ER/ERES localization until addition of biotin; after biotin treatment, the bulk of the reporter protein was found in the *cis*-Golgi, strongly overlapping with M localization, and little to no distal puncta obscured, suggesting that the reporter protein could not advance further in the secretory pathway.

**Figure 6.**
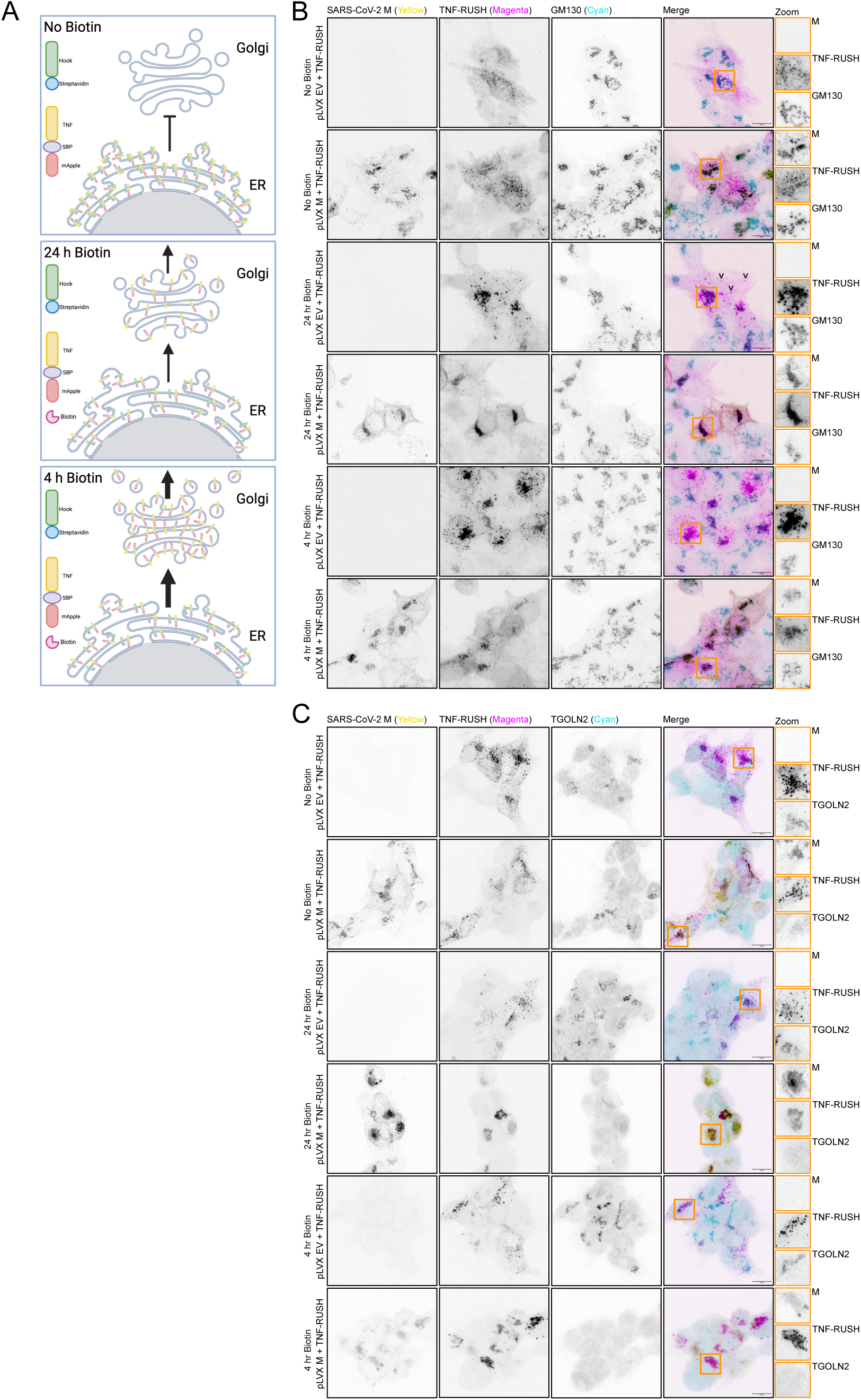
SARS-CoV-2 M inhibits protein trafficking through the secretory pathway resulting in host protein accumulation at the *cis-*Golgi. (A) Model of Retention Using Selective Hooks (RUSH) system. TNF-mApple with a strepavidin-binding peptide (SBP) is held in the ER by ER-retained streptavidin. Biotin treatment out-competes SBP-strepavidin interaction and allows TNF-mApple to continue anterograde traffic through the remained of the secretory system. **(B)** HEK293T cells were co-transfected with mApple TNF-a-RUSH with either M or EV control. Cells were then treated with 50 µM of biotin for 24h, or 4h as indicated. Cells were then fixed 24h post-transfection and stained with antibodies targeting SARS-CoV-2 M, GM130 (*cis*-Golgi), or TGOLN2 (*trans*-Golgi) and imaged by confocal microscopy. Maximum intensity projections are presented. 100X magnification, scale bar = 10 µm, Orange boxes indicate zoomed field of view. Representative images of three independent experiments.

We further leveraged our RUSH system to investigate protein trafficking in the distal secretory pathway. In the absence of M, we observed co-localization of the TNF-RUSH reporter protein with TGOLN2 after 24 h and 4 h biotin treatment, indicating proper trafficking through the secretory pathway (**Figure 6C**); however, in the presence of M we found no co-localization of our reporter protein with the TGN marker and once again the TGN was found to be dispersed in M expressing cells. Thus, SARS-CoV-2 M causes TGN dispersal, which could result from failure of cargo to escape the *cis*-Golgi.

### The M Protein Leads to Cholesterol Accumulation at the *cis*-Golgi

Efficient protein trafficking in the Golgi relies on proper distribution of cholesterol and sphingolipids in Golgi membranes. The COPI coatamer complex that mediates intra-Golgi transport and Golgi-to-ER retrograde transport is recruited to Golgi membranes by Arf GTP-binding proteins and orchestrates vesicle formation, cargo sorting, vesicle scission, and vesicle uncoating^69^. In the presence of Arf1-GTP, COPI coatamer complexes partition into liquid-disordered domains and are excluded from liquid-ordered domains rich in sphingolipids and cholesterol, resulting in the formation of vesicles bearing far fewer of these lipids than the Golgi compartments from which they derive^70^. Accordingly, these COPI-mediated transport mechanisms are sensitive to perturbations in levels of sphingolipids and cholesterol in the Golgi^71^. Considering changes in lipid and cholesterol levels in the Golgi can impair COPI-mediated transport, we hypothesized that M leads to increased levels of cholesterol at the *cis*-Golgi. To determine whether M is present in liquid-ordered domains, we isolated detergent resistant membranes (DRMs) from cells transfected with EV or M and conducted immunoblotting for cellular targets. M was found to be present in both the top (DRMs) and bottom (detergent soluble membranes (DSMs) fractions (**Figure 7A**). Calnexin was selected as a marker for ER membranes and Caveolin was selected as a positive control marker for DRMs, these markers were isolated in DSMs and DRMs, respectively. This determined that a population of M is found in the same fraction as Caveolin, confirming the presence of M in DRMs. ERGIC53 was included as marker for the ERGIC to determine whether M altered the lipid composition of the membrane for of this protein and we found ERGIC53 was isolated in DSMs in both EV and M conditions. Therefore, we confirmed M is present in liquid-ordered domains but does not alter the lipid composition of membranes containing ERGIC53.

**Figure 7.**
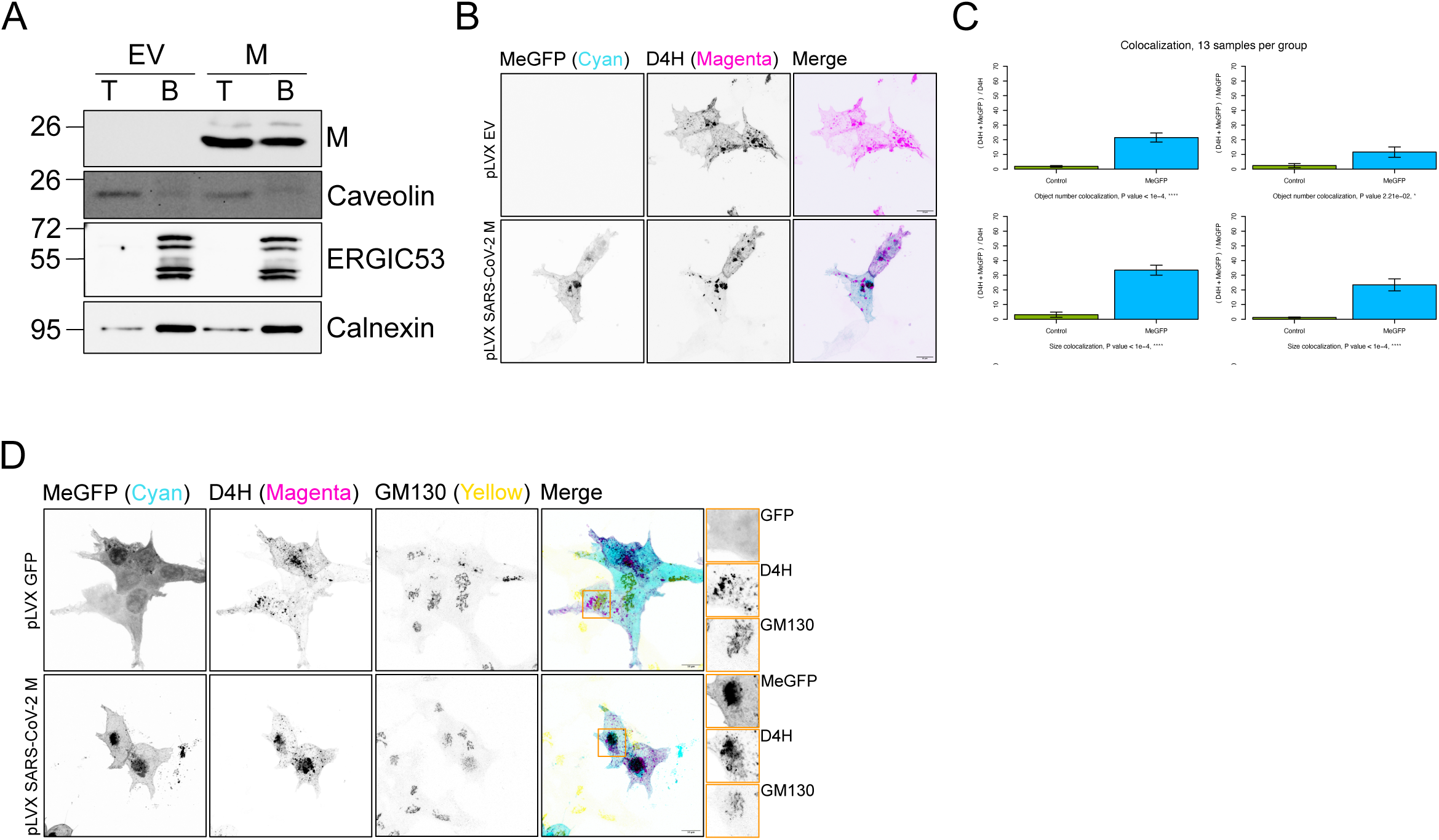
The SARS-CoV-2 M protein is present in detergent resistant membranes and increases cholesterol levels in the *cis*-Golgi. (A) HEK293T cells were transfected with M or EV and DRMs were isolated as described in methods. The top fraction, containing the DRMs and the bottom fractions were used for western blot analysis as indicated. A representative image of n = 3 is shown. **(B)** HEK293T cells were co-transfected with cholesterol biosensor D4H-mcherry and M-eGFP or EV. Cells were fixed at 24h and imaged without detergent permeabilization. **(C)** Images from 3 independent experiments as shown in **(B)** were processed using the Fiji Squassh plugin to measure object colocalization between MeGFP and D4H channels. (n = 3 ±SEM, statistical significance was determined by one-way ANOVA with Tukey-Kramer test to compare control and MeGFP groups) **(D)** HEK293T cells were co-transfected with cholesterol biosensor D4H-mcherry and M-eGFP or eGFP. Cells were fixed after 24h, and immunostained using GM-130 without permeabilizing the cells (fixation was sufficient for antibody labelling for this epitope). 100X magnification, scale bar = 10 µm, Orange boxes indicate zoomed field of view. Representative images of three independent experiments.

We next sought to determine where intracellular cholesterol localized in the presence of M. We used the D4H-mCherry cholesterol probe, which is a plasmid encoding the domain 4 (D4) of theta-toxin produced by *Clostridium perfringens* with a D434S substitution (D4H) that lowers the threshold for binding cholesterol, to label intracellular cholesterol in organelles^72–75^. HEK293T cells were co-transfected with D4H-mCherry and EV or M-eGFP, we observed that cholesterol distribution in EV cells resulted in small diffuse puncta with some larger aggregates also detected (**Figure 7B**). In cells expressing M-eGFP, cholesterol was found to be less diffuse and co-localized with M-eGFP. Using the Squassh object-based segmentation tool to measure co-localization, we found cholesterol significantly co-localized with MeGFP compared to scrambled negative control images, generated by rotating the D4H-mCherry channel to generate random co-localization^76,77^. To further confirm that co-localization of cholesterol with MeGFP was occurring at the *cis*-Golgi we conducted the same experiment and stained for the *cis*-Golgi using GM130 as a marker. In the EV condition cholesterol did not accumulate or co-localize at the *cis*-Golgi; however, in the presence of M-eGFP, cholesterol co-localized both with M and the *cis*-Golgi (**Figure 7D**). This confirms that M causes relocalization and accumulation of cholesterol at the *cis*-Golgi, likely increasing the presence of liquid ordered domains at this site.

## DISCUSSION

CoVs encode proteins that limit host antiviral responses by preventing the detection of viral products by host sensors, and if viral infection is detected, additional viral proteins prevent the synthesis of host antiviral proteins. Here, we describe another reinforcing layer of viral manipulation focused on the host secretory pathway. During infection, transmembrane CoV non-structural proteins accumulate in the ER and sculpt portions of the it into double-membrane ROs where viral RNA synthesis takes place. This is followed by a burst of viral transmembrane structural proteins including Spike, M, and E, that are synthesized and processed at the ER before proceeding to later compartments in the secretory pathway. We reasoned that these dramatic ER perturbations could affect ER proteostasis and the UPR. We observed UPR modulation by numerous CoV proteins, including Spike, which broadly activated all three arms of the UPR. By contrast, M selectively inhibited the ATF6 arm of the UPR by preventing cleavage and release of the ATF6-N transcription factor from the Golgi. M inhibited several additional processes that require ER-to-Golgi transport, including SREBP2-mediated activation of sterol responses and STING-mediated activation of interferon responses. Using a RUSH cargo sorting assay, we observed that M accumulated in the cis-Golgi and inhibited further anterograde transport of a transmembrane reporter protein beyond this compartment, while dispersing the TGN, a phenotype that we also observed during infection with diverse CoVs including SARS-CoV-2, hCoV-OC43, or hCoV-229E. Together, these observations suggest that CoV M proteins disrupt the TGN and impede normal anterograde traffic in the canonical secretory pathway. Because CoV egress does not require the TGN, this mechanism could allow the virus to selectively interfere with host responses to infection without impeding egress of nascent virions.

Golgi dispersal during viral infection is commonplace, but precise mechanisms and biological significance are not always clear^78^. Our observations of Golgi dispersal during hCoV infection is consistent with several previous reports^63–66^. One group identified multiple SARS-CoV-2 proteins that are sufficient to trigger Golgi dispersal, including M^64^. However, in this study M was found to overlap with the ER marker calnexin, and the authors suggested that membranous structures formed from ectopic expression of M resembled smaller double membrane vesicles (DMVs). By contrast, using immunostaining and a cargo sorting assay, we observed accumulation of M at the *cis*-Golgi, along with retention of diverse cargo proteins at this compartment. Our observations are consistent with accumulating evidence that Golgi dispersal during viral infection impedes protein trafficking in the secretory pathway.

How does M impede protein cargo trafficking in the secretory pathway, and could it relate to its role in virion assembly? M is a polytopic transmembrane glycoprotein, with a short lumenal N-glycosylated amino-terminal domain, three transmembrane helices connected by short loops, and a cytoplasmic hinge region that connects to an extended carboxy-terminal β-sheet sandwich domain (BD)^35^. M is the most abundant viral structural protein, and virion assembly places the short amino terminal domain on the exterior of the virion, the transmembrane domains in the virion envelope, and the extended carboxy-terminal domain in the virion interior, where it interacts with the viral genome. Structural studies have revealed the importance of M dimer formation and the formation of higher-order oligomers in the virus assembly process^35,79,80^. Cryo-electron microscopy studies demonstrated that M forms homodimers where three-helix transmembrane domain bundles are comprised of helices from different monomers (transmembrane domain 1 assembles with domains 2 and 3 from the binding partner), and lateral interactions of these dimers in the membrane can lead to tetrameric and hexameric oligomers. Furthermore, these M dimers exist in two forms dictated by conformational changes in the hinge region, a “short” form with a compact appearance, and a “long” form with an extended appearance. Both forms of M are required for virus assembly and can be found in intact virions^35^, with the short form associated with membrane flexibility and low Spike density, and the long form associated with more rigid membranes and Spike clusters^79^. These two forms of M exist in dynamic equilibrium, and regulated transition between these states is required to drive virion assembly. This model is supported by the recent discovery of two small molecule inhibitors of CoV assembly that bind the hinge region of M and either stabilize the short form^81^ or stabilize a transition intermediate between the short and long forms^82^. Thus, the dynamic toggling of M between these two forms, and its influence on local membrane properties, appears to be critical for virion assembly. Atomic force microscopy studies have shown M dimers associated with ‘thin’ membranes^83^, and others have shown that M directly interacts with sphingolipids including ceramide-1-phosphate (C1P) in a conformationally selective manner^84^, with molecular dynamics simulations revealing stabilization of the short form of M. Ceramide kinase (CERK) is a *cis*-Golgi-resident enzyme that converts ceramide into C1P, providing an attractive mechanism for accumulation of M in this compartment.

In a recent study of global subcellular protein reorganization during hCoV-OC43 infection, all seven protein subunits of the COPI complex were mislocalized, which was confirmed by demonstrating that COPE and COPB2 redistribute from a perinuclear location to a broadly dispersed cytoplasmic pattern during infection^85^. Since COPI is responsible for intra-Golgi trafficking, and we have shown that M causes accumulation of protein cargo and cholesterol at the *cis*-Golgi, we speculate that M could increase liquid-ordered domains at the *cis*-Golgi that could support virus assembly at the expense of COPI recruitment to these sites, potentially impairing Golgi protein trafficking and triggering TGN dispersal. This notion of M supporting the formation of rigid ordered lipid domains at sites of CoV assembly is reminiscent of the reports that Spike palmitoylation aids the formation of ordered lipid nanodomains enriched in cholesterol and sphingolipids at ERGIC membranes that aid proper assembly of infectious virions^86^. Importantly, preventing Spike palmitoylation by substituting key cytoplasmic cysteine residues prevented Spike from accumulating at these detergent-resistant membranes, but M persisted, suggesting an independent mechanism for M recruitment to these membranes.

Taken together, our findings provide important new information about how a viral structural protein can moonlight as a host shutoff protein by interfering with subcellular protein trafficking, possibly as a result of its role in nucleating sites of virion assembly in the early secretory pathway. Considering the newfound focus on M as a target for antiviral small molecules, our study provides a pathway for better understanding the dynamic interactions of M and other viral structural proteins, along with key host sterols and lipids, at these sites of assembly.

## MATERIALS AND METHODS

### Cells and viruses

Human embryonic kidney (HEK) 293T, human adenocarcinoma alveolar basal epithelial A549-ACE2 and human hepatoma Huh7.5 cells were grown in Dulbecco’s modified Eagle’s medium (DMEM; Thermo Fisher, 11965118) supplemented with heat-inactivated 10% fetal bovine serum (FBS, Thermo Fisher, A31607-01), 100 U/mL penicillin, 100 µg/mL streptomycin, and 2 mM L-glutamine (Pen/Strep/Gln; Thermo Fisher, 15140122 and 25030081). All cells were maintained at 37°C in a 5% CO_2_ atmosphere. To generate 293T-GFP-ERGIC53 cells, 293T cells were stably transduced with a lentivirus vector encoding GFP-ERGIC53 (pLVX-GFP-ERGIC53-Puro, Addgene #134859), then selected and maintained in 10 µg/mL Puromycin (Gibco, A1113803). A549-ACE2 cells were generated by lentivirus transduction of ACE2 as in^87^. Cells were maintained for 48-72 hours in 5 µg/mL blasticidin (Thermo Fisher, A1113903) to select for ACE2-expressing cells. Following selection, cells were cultured in DMEM with 10% FBS and Pen/Strep/Gln.

All experiments with severe acute respiratory syndrome coronavirus 2 (SARS-CoV-2) were conducted in the University of Calgary Containment-level 3 (CL3) facility in accordance with the CL3 Oversight Committee and Biosafety Office regulations. SARS-CoV-2 Toronto-1 variant was propagated in Vero-E6 cells, as in^88^. Briefly, cells were infected at a MOI of 0.01 for 1 h in serum-free DMEM. Following adsorption, cells were maintained in DMEM supplemented with 2% FBS and Pen/Step/Gln at 37°C. Five days post-infection, virus-containing media was centrifuged at 1000 x g for 5 min, aliquoted, and stored at -80°C. SARS-CoV-2 was not passaged beyond passage three. SARS-CoV-2 titers were determined by plaque assay in Vero-E6 cells as in^88^ using equal parts 2.4% colloidal cellulose (Sigma, cat # 425244; prepared in sterile H2O) and 2X DMEM (Wisent) supplemented with 1% FBS and Pen/Strep/Gln. 72 hours post infection, cells were fixed, stained, and plaques were enumerated.

Stocks of human coronavirus 229E (HCoV-229E; ATCC, VR-740) were propagated in Huh7.5 cells. Cells were infected at a MOI of 0.05 for 1 h in serum-free DMEM. After 1 h, the infected cells were maintained in DMEM supplemented with 2.5% FBS and Pen/Strep/Gln for five days at 33°C. Upon harvest, the culture supernatant was centrifuged at 1000 x g for 5 min at 4°C, aliquoted, and stored at -80°C. Stocks of recombinant HCoV-OC43-mClover^89^ were propagated in BHK-21 cells. Cells were infected at a MOI of 0.05 for 1 h at 37°C in serum-free DMEM. After 1 h, the infected cells were maintained in DMEM supplemented with 1% FBS and Pen/Strep/Gln until cytopathic effect was complete. Upon harvest, the culture supernatant was centrifuged at 1000 x g for 5 min at 4°C, aliquoted, and stored at -80°C. Viral titers were measured using median tissue culture infectious dose (TCID50) assays using the Spearman-Kärber method.

. Following the serial dilutions of the samples, the appropriate cell line was infected for 1 h at 37°C prior to the replacement of the inoculum for indicated overlay medium and incubated at 37°C: Huh7.5 cells were infected with hCoV-229E in DMEM/2.5%FBS/Pen/Strep/Gln medium, while BHK-21 cells were infected with recombinant hCoV-OC43-mClover in DMEM/1%FBS/Pen/Strep/Gln medium.

To generate lentiviruses for stable transductions, 293T cells were transfected in a 10 cm dish with 3 µg pLVX-GFP-ERGIC-Puro (Addgene #134859) + 2 µg psPAX2 (a gift from Didier Trono; Addgene #12260) + 1 µg pMD2.G (a gift from Didier Trono; Addgene #12259) and 18 µL PEI diluted in Opti-MEM (Gibco, 31985070). After 4 h, the medium was changed to 293T growth medium. After 48 h, supernatants were harvested, cleared by filtration using a 0.45 µm filter, aliquoted, and stored at -80°C until use.

### Chemicals

Tunicamycin (Tm, T7765), Thapsigargin (Tg, T9033), Brefeldin A (BFA, B7651), Cerivastatin (Statin, SML0005), and Biotin (B4501) were purchased from Sigma-Aldrich. Tm, Tg, BFA and biotin were solubilized in dimethyl sulfoxide (DMSO) and Statin was solubilized in water. All drugs were stored at -80°C. Stock concentrations were diluted to the indicated concentrations in cell culture media.

### Plasmids

All plasmids were purified using the QIAprep Spin Miniprep or QIAfilter Plasmid Midi kits (QIAGEN) and all restriction enzymes were purchased from New England Biolabs (NEB). All SARS-CoV-2 2xStrep tagged constructs used in the ORF library screen (**Figure 1**) were from the pLVX-SARS-CoV-2 ORF library^43^ (kind gifts from Nevan Krogan). The following plasmids were purchased from Addgene: : pLDLR-luc (aka: pES7) (#14940, a kind gift from Axel Nohturfft), pTRIP-CMV-tagRFP-FLAG-cGAS and pMSCV-hygro-STING (#86676, #102598, kind gifts from Nicolas Manel), mApple-TNFa-RUSH (#166902, a kind gift from Jennifer Lippincott-Schwartz), pLVX-EF1a-EGFP-ERGIC53-IRES-Puromycin (#134859, a kind gift from David Andrews), and lenti-CRISPRv2 (#52961, a kind gift from Feng Zhang). The pCMV-*Gaussia* Luc Vector was purchased from ThermoFisher (16147). The pLJM1-Luc2 vector was generated by cloning Luc2 from pGL4.26 (Promega) into pLJM1-B*-Puro^90,91^. The pcDNA3.1+-SARS-CoV-2-Spike (D614) plasmid contains a codon-optimized ORF for Spike from GenBank NC_045512 that was synthesized by GenScript (a kind gift from David Kelvin) then cloned between the *Kpn*I and *Bam*HI sites of pcDNA3.1(+)^90^. To generate pLVX M no tag, M was PCR amplified from the pLVX-M-2xStrep construct from the Krogan library and cloned back into pLVX with *Eco*RI and *Bam*HI. To generate pLVX E no tag, E was PCR amplified from the pLVX-M-2xStrep construct from the Krogan library and cloned back into pLVX with *Eco*RI and *Bam*HI. All the additional hCoV M proteins (SARS1, MERS, hCoV-OC43, hCoV-HKU1, hCoV-229E, and hCoV-NL63) were synthesized by GenScript in pcDNA3.1 (-) using the GenBank sequences NC_004718.3, NC_019843.3, AY391777.1, NC_006577.2, NC002645.1 and NC_005831.2, respectively. All hCoV M sequences were then cloned into pLVX using *Eco*RI and *Bam*HI. To generate pLJM1-B*-Puro-HA-ATF6^91^ and pLJM1-B*-Puro-HA-ATF6-N, HA-ATF6 and HA-ATF6-N were PCR amplified from pCGN-ATF6 and pCGN-ATF6 (1-373) (Addgene, #11974 and #27173, kind gifts from Ron Prywes) and cloned into pLJM1-B*-Puro with *Nhe*I and *Age*I. To generate the ERSE-luciferase construct pairs of phosphorylated and annealed oligos for 2 ERSE sequences separated by a spacer, were cloned into pGL4.26 (Promega) using *Bgl*II and *Acc*65I. To generate the CMV-Renilla construct the Renilla gene was cloned from pcDNA3-RLUC-POLIRES-FLUC (Addgene, #45642, a kind gift from Nahum Sonenberg) into pcDNA3.1(+) using *Nhe*I and *Kpn*I. To generate the pLVX-EV construct pairs of phosphorylated and annealed oligos were cloned into pLVX M no tag, cutting out the M gene with *Eco*RI and *Bam*HI. The pGL4.26-ISRE (ISRE-luciferase) construct was generated by cloning the 5xISRE response elements from pISRE-luc (Stratagene) with *Bam*HI and *Eco*RI (blunted with Klenow) into pGL4.26 (Promega) with *Bgl*II and *HinD*III (blunted with Klenow). The pGL4.26-NFkB (NFkB-luciferase) construct was generated by cloning the 4x NFkB response elements from pNFKB-luc (Stratagene) into pGL4.26 (Promega) using *Nhe*I and *Bgl*II. Primer sequences for PCR amplification and annealed oligos can be found in Table 1.

**Table 1.**
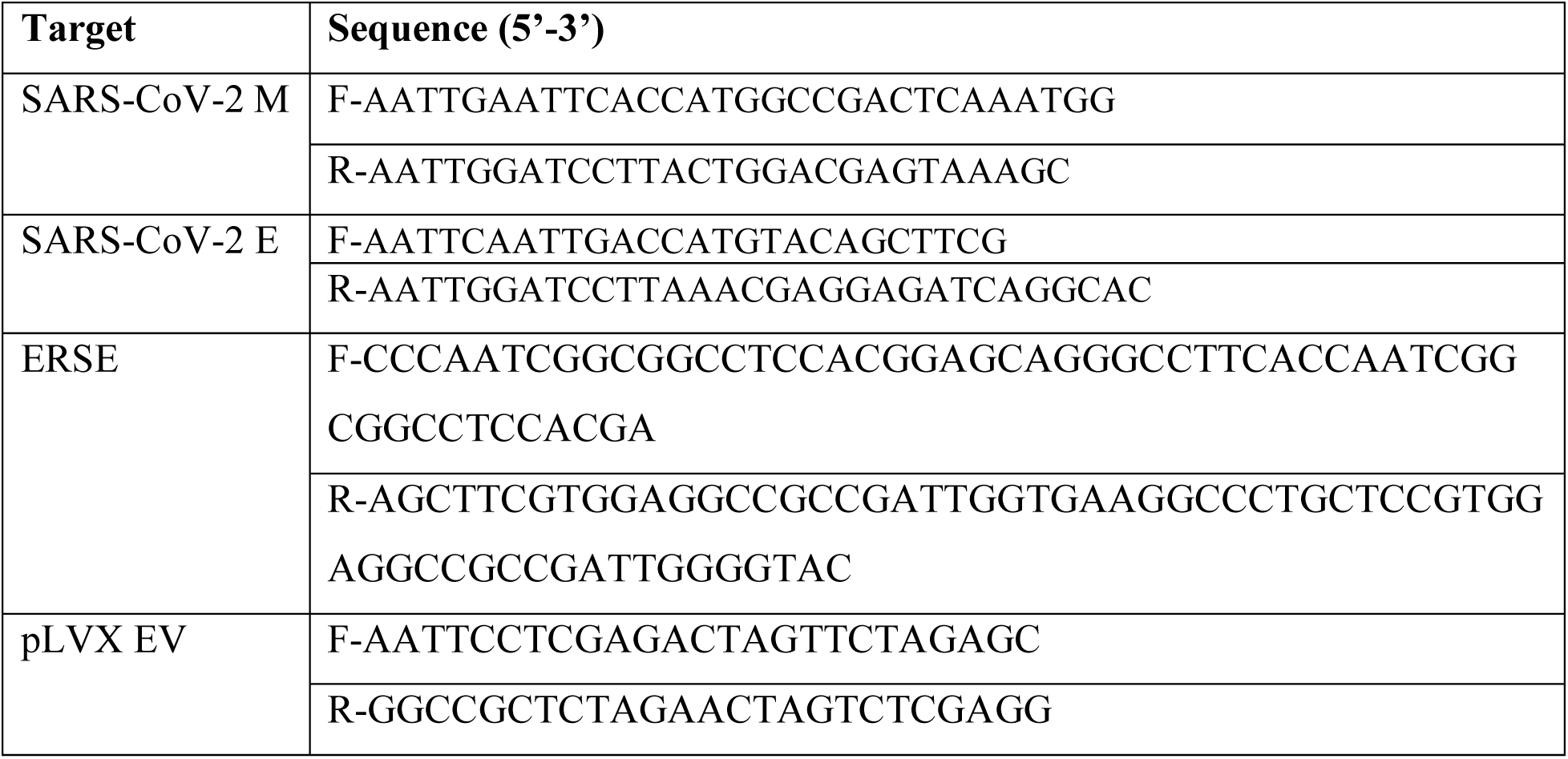
Primers used for cloning.

### Transfections

HEK293T cells were transfected using polyethylenimine (PEI); Linear, MW 40000 (Polyscience, 24765) dissolved in water (pH 7.4). Both plasmids and PEI were diluted in Opti-MEM I (Thermo Fisher, 31985070) and incubated for 5 min, then combined and incubated for 15 min before adding to cell monolayers in antibiotic free 10%FBS/Gln DMEM. For drug treatments, drugs were added to media containing the transfection reaction at indicated times without a media change, excluding the Gaussia-luciferase assays in which media was removed at 24 h post-transfection and replaced with fresh media and BFA for indicated samples.

### CRISPR/Cas9-mediated deletion of human ATF6

HEK293T cells were seeded in a 6-well plate. The following day cells were stably transduced with lentiviruses containing pLentiCRISPRv2-ATF6 plasmids. At 24 h post-transduction cells were selected and maintained in 10 µg/mL Puromycin (Gibco, A1113803). After expansion of cells under selection, monoclonal ATF6 knockout populations were generated by seeding cells into a 96-well plate at 0.5 cells per well. Wells were screened to ensure populations were expanded from single cells. Individual monoclonal populations were expanded in 6-well plates and screened via immunoblotting for successful deletion of ATF6. A single clone was selected and used throughout the study.

To generate lentiviruses for stable transductions, HEK293T cells were transfected in a 10 cm dish with 3 µg pLentiCRISPRv2-ATF6 + 2 µg psPAX2 (a gift from Didier Trono; Addgene #12260) + 2 µg pMD2.G (a gift from Didier Trono; Addgene #12259) and 18 µL PEI diluted in Opti-MEM (Gibco, 31985070). After 4 h, the medium was changed to HEK293T growth medium. After 48 h, supernatants were harvested, cleared by filtration using a 0.45 µm filter, aliquoted, and stored at -80°C until use.

To generate pLentiCRISPRv2-ATF6 annealed oligos targeting sequences within the ATF6 gene were cloned into pLentiCRISPRv2 using the following sequences: F-5’-CACCGTTTGCCAATGGCATAAGCGT-3’ and R-5’-ACGCTTATGCCATTGGCAAACGGTG-3’.

### Luciferase reporter assays

HEK293T cells were seeded in 96-well plates coated with poly-L-lysine (Sigma, P2658). The following day cells were transfected with plasmids as indicated in figures. At 24 h post-transfection cells were lysed in 1x Reporter Lysis Buffer (Promega, E397A). For Gaussia-luciferase experiments media was changed at 24 h post-transfection and at 30 h post-transfection supernatant was collected for analysis and cells were lysed as previously indicated. Lysates and supernatant were stored at -80°C until analysis. Once thawed, 10 µL of lysate or supernatant was added to a white 96-well plate (Costar, 3917). Measurements of firefly and Renilla luciferase were conducted using the Promega Dual Luciferase Kit (Promega, E1910 or E1960), substrates and buffers were prepared as indicated by the manufacturer. Plates were read on a CLARIOstar^PLUS^ (BMG-Labtech, Serial #430-2826, software version 5.70 R3) and raw data readings were collected in Data Analysis Mars (software version 4.00 R2). pLVX empty vector (EV) or pcDNA3.1(+) were used as controls as indicated for normalization of proteins expressed in the respective vectors.

### Semi-quantitative *XBP1* splicing assay

HEK293T cells were seeded in 12-well plates coated with poly-L-lysine (Sigma-Aldrich, P2658). The following day cells were transfected with pcDNA3.1, pcDNA3.1-SARS-CoV-2-Spike-D614, and pLVX-SARS-CoV-2-M, as indicated. Total RNA from cells was extracted using the RNeasy Plus Mini Kit (QIAGEN, 74134) following the manufacturer’s protocol. Synthesis of cDNA was performed using the Maxima H Minus First Strand cDNA Synthesis Kit (ThermoFisher, K1652) using random hexamer primers. A 473-bp PCR product spanning exon/intron boundaries was generated using the *XBP1* forward primer 5’-AAACAGAGTAGCAGCTCAGACTGC-3’ and the *XBP1* reverse primer 5’-TCCTTCTGGGTAGACCTCTGGGAG-3’. The PCR product was digested overnight with PstI-HF to cleave the unspliced *XBP1* product into XBP1u1 and XBP1u2. The digested PCR product was resolved on a 2.5% agarose gel made with 1x Tris-acetate-EDTA and stained with ethidium bromide (Sigma-Aldrich, E1510). The gel was imaged using a ChemiDoc MP Imaging system (Bio-Rad).

### Immunoblotting

Cell monolayers were washed once with PBS and lysed in 2x Laemmli buffer (4% [wt/vol] sodium dodecyl sulfate (SDS), 20% [vol/vol] glycerol, 120 mM Tris-HCl [pH 6.8]). DNA was sheared by repeated pipetting with a 21-gauge needle before adding 100 mM dithiothreitol (DTT), bromophenol blue, and boiling at 95°C for 5 min. Samples were stored at -20°C until analysis. Total protein concentration was determined by DC protein assay (Bio-Rad, 5000116) against a bovine serum albumin (BSA) standard curve and measured in a 96-well plate format at 750 nm using an Eon (BioTek) microplate spectrophotometer. Equal quantities of 10 µg total protein were loaded in each SDS-PAGE gel, with Color Prestained Protein Standard, Broad Range (NEB, P7719S), and separated at 100 V. Proteins were transferred to polyvinylidene fluoride (PVDF) membranes using the Trans-Blot Turbo RTA Midi 0.2 µm PVDF Transfer Kit (Bio-Rad, 1704273) and a Trans-Blot Turbo Transfer System (Bio-Rad). Membranes were blocked with 5% BSA or 5% skim milk (PERK, BiP, and ATF6 blots) in tris-buffered saline/0.1% [vol/vol] tween-20 (TBS-T) before probing overnight at 4°C with antibodies in 5% BSA in TBS-T raised to the following targets: rabbit anti-SARS-CoV-2 S1 RBD (Elabscience, E-AB-V1006, 1:2000), rabbit anti-SARS-CoV-2 M (Novus Biologicals, NBP3-05698, 1:2000), rabbit anti-SARS-CoV-2 E (abbexa, abx226552, 1:2000), rabbit anti-PERK (Cell Signaling Technologies (CST), #5683, 1:2000), mouse anti-CHOP (CST, #2895, 1:1000), mouse anti-XBP1s (CST, #12782, 1:1000), rabbit anti-BiP (CST, #3177, 1:1000), mouse anti-HA (CST, #2367, 1:1000), mouse anti-ATF6 (AbCam, ab122897, 1:1000), rabbit polyclonal anti-Caveolin (BD Biosciences, C13630), rabbit anti-ERGIC-53 (Sigma, E1031), and rabbit anti-β-actin (CST, #8457, 1:1000). Membranes were washed with TBS-T and incubated with HRP-linked secondary antibodies for 1 h at room temperature in 5% BSA in TBS-T. Secondary antibodies used: anti-rabbit, HRP-linked (CST, #7074, 1:3000), and anti-mouse, HRP-linked (CST, #7076, 1:3000). Blot were developed with Clarity ECL chemiluminescence reagent (Bio-Rad, 170-5061) or Clarity Max ECL chemiluminescence reagent (Bio-Rad, 170-5062) for PERK, ATF6 and BiP blots. All blots were imaged using a ChemiDoc MP Imaging System (Bio-Rad). Molecular weights were determined using the Broad Range, Color Prestained Protein Standard (NEB, P7719S). Molecular weights in kDa are indicated on the left of blot images. Images were analyzed using Image Lab 6.1 (Bio-Rad). Images were cropped and annotated using Affinity Designer (Serif).

### Immunofluorescence microscopy

HEK293T cells were seeded on #1.5 coverslips (Paul Marienfeld GmbH & Co. KG, 0117580) coated with poly-L-lysine (Sigma-Aldrich, P2658). The following day cells were transfected or infected with indicated plasmids or viruses. To harvest, the cells were washed once with PBS and fixed with 4% paraformaldehyde (Electron Microscopy Services, 15710) in PBS for 15 min at room temperature. Coverslips were blocked and permeabilized in staining buffer (1% human serum (Sigma-Aldrich, 4552) heat-inactivated at 56°C for 1 h, 0.1% Triton X-100 in PBS) for 1 h at room temperature. Coverslips were stained overnight at 4°C with the following antibodies as indicated: mouse anti-SARS-CoV-2 M (R&D Systems, MAB10696), rabbit anti-GM130 (CST, 12480S, 1:3000), rabbit anti-TGOLN2 (BETHYL, A304-434A, 1:200) and mouse anti-dsRNA J2 (SCICONS, RNT-SCI-10010200). The following day the coverslips were washed three times with PBS, then stained with secondary antibodies: goat anti-mouse-555 (Invitrogen, A21422, 1:1000), anti-mouse-647 (Invitrogen, A21463, 1:1000) goat anti-rabbit-647 (Invitrogen, A21244, 1:1000), anti-rabbit-488 (Invitrogen, A21441, 1:1000) donkey anti-rabbit-555 (Invitrogen, A31572, 1:1000), and chicken anti-mouse-488 (Invitrogen, A21200, 1:1000) in staining buffer for 1 h at room temperature in the dark. Coverslips were washed with PBS three times, and then counterstained for 5 min with Hoescht 33342 (ThermoFisher, 62249). Coverslips were then mounted on cover glass (Fisher Scientific, 12-550)) using ProLong Gold anti-fade reagent (ThermoFisher, P36930). Z-stacks were imaged on a Zeiss LSM880 and processed into maximum intensity projections using Zen Black (Zeiss).

### Retention Using Selective Hooks (RUSH) cargo sorting assay

HEK293T cells were transfected with mApple-TNFa-RUSH, along with pLVX EV, or pLVX M as indicated. Cells were treated with biotin (Sigma, B4501) at time of transfection (24 h treatment) or at 20 hpt. At 24 hpt cells were fixed, stained and imaged as described above.

### Isolation of detergent resistant membranes (DRM) from cells

DRMs were isolated essentially as described in [PMID: 17947970]. Breifly, HEK293T cells were seeded into 3-wells of a 6-well cluster dish and tranfected with pLVX-M or EV the following day as described above. Cells were then washed once with ice-cold PBS, then once-with TNE buffer (150 mM NaCl, 2 mM EDTA, 50 mM Tris–HCl, pH 7.4) then scraped and washed once more in TNE buffer. The cell pellet was then resuspended in 200 µL TNE with cOmplete EDTA-free Protease Inhibitor Cocktail (Roche) then homogenized with 25-G needle (25-strokes). 180 µL of the lysed cell suspension was transferred to a new tube with 20 µL of freshly prepared 10% Triton X-100, mixed by inversion, and incubated on ice for 30 min. Then 400 µL of Optiprep (60% iodixanol; Sigma) added, mixed, and the resulting 600 µL was moved to an ultracentrifuge tube for a S-55S (Thermo) ultracentrifuge rotor and carefully overlaid with 1.2mL 30% iodixanol-TNE then 300 µL of TNE without iodixanol. The tubes were then transferred to the S-55S rotor and centrifuged at 55,000 rpm for 2h. Two 1 mL fractions were then removed from the tube and used for immunoblot analysis as described above.

### Data management and analysis

Graphing and statistical calculations were performed using GraphPad Prism for macOS v10.3.1. Figures were prepared using Affinity Designer v1.10.8 (Serif). Raw microscopy images were processed and intensities standardized using ImageJ2 v2.16.0/1.54n. Merged microscopy images were uploaded to https://amsterdamstudygroup.shinyapps.io/ezreverse/ [https://doi.org/10.1101/2024.05.27.594095] for image inversion.

## ACKNOWLEDGEMENTS

We thank the following colleagues for their assistance this work: Dr. Roy Duncan and Nichole McMullen for the use of the CLARIOstar^PLUS^ plate reader. We thank Dalhousie University Core Facility managers Dr. Gerard Gaspard (Cellular Molecular Digital Imaging) and Dr. Christopher Hughes (Biological Mass Spectrometry) for expert technical support. We thank Dr. Nevan Krogan (UCSF) for the generous gift of a collection of lentiviral vectors expressing SARS-CoV-2 ORFs^43^. The following plasmids were gifts provided through Addgene: pLVX-EF1a-EGFP-ERGIC53-IRES-Puromycin (David Andrews, https://www.addgene.org/134859/), mApple-TNFa-RUSH (Jennifer Lippincott-Schwartz, https://www.addgene.org/166902/); pLDLR-Luc (aka: pES7) (Axel Nohturfft, https://www.addgene.org/14940/); pTRIP-CMV-tagRFP-FLAG-cGAS and pMSCV-hygro-STING (Nicola Manel, https://www.addgene.org/86676/, https://www.addgene.org/102598/); lenti-CRISPRv2 (Feng Zhang https://www.addgene.org/52961/). This work was supported by Canadian Institutes for Health Research (CIHR; https://cihr-irsc.gc.ca) Project Grant PJT-148727 (to C.M.), Coronavirus Variants Rapid Response Network (CoVaRR-Net; https://covarrnet.ca) Grant 175622 (to J.A.C. and others), and Nova Scotia COVID-19 Health Research Coalition Grants (https://researchns.ca/covid19-health-research-coalition/) to C.M., and E.S.P. The funders had no role in study design, data collection and analysis, decision to publish, or preparation of the manuscript.

## AUTHOR CONTRIBUTIONS

Conceptualization: T.C., E.S.P., C.M.; Methodology: T.C., E.S.P., C.M.; Formal Analysis: T.C.,

E.S.P.; Investigation: T.C., E.S.P., R.P.M.; Resources: T.C., E.S.P., R.P.M.; Writing – Original

Draft: T.C., E.S.P., C.M.; Writing –Review & Editing: T.C., E.S.P., R.P.M., J.A.C., C.M.;

Visualization: T.C., E.S.P.; Supervision: E.S.P., J.A.C., C.M.; Project Administration: J.A.C., C.M.; Funding Acquisition: J.A.C., C.M.

## DECLARATION OF INTERESTS

All authors declare no conflicts of interest.

**Supplementary Figure 1.** S**u**ccessful **knockout of ATF6 from HEK293T cells.** HEK293T cells were transduced with lentiviruses encoding pLentiCRIPSRv2-ATF6 constructs to knockout the ATF6 gene. At 24 h post-transduction cells were selected in 10 mg/mL of puromycin. Cells were seeded in a 96-well plate to generate monoclonal populations and expanded under puromycin selection. Selected monoclonal populations were seeded into 6-well plates, lysates were harvested 48 hours post-seeding and stored at -20C prior to immunoblotting. * indicates the knockout clonal population selected for further experimental use.

