## Supplementary material for "Coronavirus M proteins disperse the trans-Golgi network and inhibit anterograde protein trafficking in the secretory pathway": Figure S1

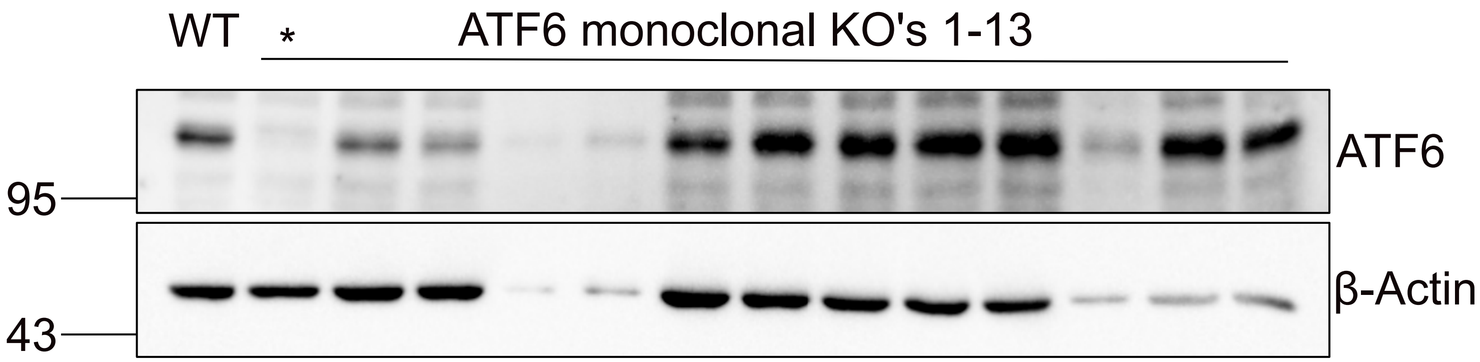


**Supplementary Figure 1. Successful knockout of ATF6 from HEK293T cells.** HEK293T cells were transduced with lentiviruses encoding pLentiCRIPSRv2-ATF6 constructs to knockout the ATF6 gene. At 24 h post-transduction cells were selected in 10 mg/mL of puromycin. Cells were seeded in a 96-well plate to generate monoclonal populations and expanded under puromycin selection. Selected monoclonal populations were seeded into 6-well plates, lysates were harvested 48 hours post-seeding and stored at -20C prior to immunoblotting. * indicates the knockout clonal population selected for further experimental use.
